# DNA extraction and host depletion methods significantly impact and potentially bias bacterial detection in a biological fluid

**DOI:** 10.1101/2020.08.21.262337

**Authors:** Erika Ganda, Kristen L. Beck, Niina Haiminen, Justin D. Silverman, Ban Kawas, Brittany Cronk, Renee R. Anderson, Laura B. Goodman, Martin Wiedmann

## Abstract

Untargeted sequencing of nucleic acids present in food can inform the detection of food safety and origin, as well as product tampering and mislabeling issues. The application of such technologies to food analysis could reveal valuable insights that are simply unobtainable by targeted testing, leading to the efforts of applying such technologies in the food industry. However, before these approaches can be applied, it is imperative to verify that the most appropriate methods are used at every step of the process: gathering primary material, laboratory methods, data analysis, and interpretation.

The focus of this study is in gathering the primary material, in this case, DNA. We used bovine milk as a model to 1) evaluate commercially available kits for their ability to extract nucleic acids from inoculated bovine milk; 2) evaluate host DNA depletion methods for use with milk, and 3) develop and evaluate a selective lysis-PMA based protocol for host DNA depletion in milk.

Our results suggest that magnetic-based nucleic acid extraction methods are best for nucleic acid isolation of bovine milk. Removal of host DNA remains a challenge for untargeted sequencing of milk, highlighting that the individual matrix characteristics should always be considered in food testing. Some reported methods introduce bias against specific types of microbes, which may be particularly problematic in food safety where the detection of Gram-negative pathogens and indicators is essential. Continuous efforts are needed to develop and validate new approaches for untargeted metagenomics in samples with large amounts of DNA from a single host.

**Importance:** Tracking the bacterial communities present in our food has the potential to inform food safety and product origin. To do so, the entire genetic material present in a sample is extracted using chemical methods or commercially available kits and sequenced using next-generation platforms to provide a snapshot of what the relative composition looks like. Because the genetic material of higher organisms present in food (e.g., cow in milk or beef, wheat in flour) is around one thousand times larger than the bacterial content, challenges exist in gathering the information of interest. Additionally, specific bacterial characteristics can make them easier or harder to detect, adding another layer of complexity to this issue. In this study, we demonstrate the impact of using different methods in the ability of detecting specific bacteria and highlight the need to ensure that the most appropriate methods are being used for each particular sample.

## INTRODUCTION

In the past decade, recent developments in molecular methods, including high throughput sequencing (**HTS**) technologies, have demonstrated the feasibility of sequencing-based analysis of various foods and food-associated environments, and the potential for application in informing food safety practices (1), product processing methods (2) and ingredient authentication (3, 4). This has spurred the efforts to translate the use of such technologies to the industry setting with the objective of moving food safety testing to the next frontier (5). However, before such refined approaches can be reliably applied in the industry, it is imperative that appropriate methods are employed at every step of the process: gathering primary material, laboratory methods, data analysis, and interpretation. In this study, we highlight the importance of considering the food matrix characteristics as well as the laboratory methodologies applied. Bovine milk is used as a model with to demonstrate the several challenges associated with developing HTS methods for use in food matrices.

Milk, similar to many other foods, is a chemically complex biological fluid. It contains several compounds that can hamper the chemistry involved in DNA and RNA extraction (6, 7) and act as PCR inhibitors, such as calcium ions, fats, and proteins (8, 9). Additionally, lactoferrin, an enzyme present in bovine milk, has recently been described to have both DNase and RNase activity (10). Another challenge for untargeted sequencing applications of bovine milk is the presence of bovine somatic cells. The bovine genome is one thousand times larger than an average bacterial genome (Bovine: 2.7 Gb, Bacteria: average 3.6 Mb, median 3.4 Mb (11, 12)). Thus, even when present in much smaller amounts than bacterial cells, bovine somatic cells introduce an enormous quantity of typically unwanted host nucleic acids in untargeted HTS studies. Realistically, however, high-quality raw milk may contain around 200,000 bovine somatic cells and 20,000 or less bacterial cells per mL, leading to a 10,000-fold higher abundance of bovine DNA in high quality raw milk.

Despite the challenges associated with nucleic acid extraction from milk and the amount of host DNA present, several investigations have successfully used milk (13–15) or other dairy products (16) as their sample of interest in targeted and untargeted HTS studies, highlighting the potential for application of HTS technologies in food production settings. Total RNA sequencing can be more informative than untargeted DNA sequencing as it has the potential to provide gene expression (e.g. toxin production) in addition to taxonomic relative abundance in a community of food-associated microorganisms (4, 17). When compared to DNA extraction, RNA extraction is a more complex and challenging process, given the short half-life of RNA compared to DNA, its inherent susceptibility to degradation, and the known presence of RNases in milk (10). Nevertheless, a number of studies have been successful in extracting RNA from dairy products highlighting the potential of RNA based techniques for food safety and quality surveillance (18–20).

When the ultimate objective is to use HTS to create a tool that can be used to inform food safety and quality in industry settings, understanding the impact and biases introduced by using protocols that have not yet been tested or optimized for a given food matrix is of great importance. In addition, to be potentially adopted in industry settings, laboratory protocols should be performed in a timely manner. It is important to characterize the entire food sample, and most protocols available for nucleic acid extraction in milk begin with a centrifugation and fat-removal step, which can in itself introduce significant bias to the final result. For example, bacterial spores have been described to aggregate in the fat layer of milk samples subjected to gravity separation (21, 22), and RNA yield has been shown to be variable among different milk fractions (23). Unfortunately, these potential introductions of bias are overlooked in most investigations.

While a number of studies have investigated commercially available protocols for nucleic acid extraction of milk in terms of DNA concentration, quality, and ability to detect specific pathogens (24–26), their main objective was not the assessment of differential DNA extraction nor biases in the representation of diverse bacterial populations, which would require the inclusion of mock bacterial communities of interest into a milk sample and comparison of several protocols.

Host DNA contamination is a challenge not exclusive to milk and other food samples, as mammalian DNA has been shown to dominate the number of sequencing reads in cerebrospinal fluid, skin, vaginal, and oral metagenomes in humans(27–29). To tackle the host DNA issue, enzymatic and immunomagnetic protocols aimed at decreasing host DNA contamination became commercially available and have been tested in select sample types (27, 28, 30). In addition, a number of “homebrew” methods have been tested to allow for successful depletion of host DNA (27, 31, 32). Nevertheless, these methods are not guaranteed to work in every sample type, and to the best of our knowledge, no host DNA depletion methods have been evaluated for their applicability in milk.

The goals of this study were 1) to evaluate commercially available kits for their ability to extract nucleic acids from bovine milk; 2) to evaluate host DNA depletion methods for use with bovine milk, and 3) to develop and evaluate a selective lysis-PMA based protocol for host DNA depletion in milk. Our overarching hypothesis was that methodologies would differ regarding the efficacy of nucleic acid extraction, and potential biases would be observed. Experiments were thus performed on raw bovine milk inoculated with mock bacterial communities, which included Gram-negative (*Salmonella enterica*), Gram-positive (*Listeria monocytogenes*), *Mycobacteria* (*M. smegmatis*) as well as spores representing aerobic sporeformers (*Bacillus wiedmannii*).

## METHODS

### Comparison of Nucleic Acid Extraction Protocols in Spiked Milk

#### Strain Selection

We selected a Gram-positive (*L. monocytogenes*), a Gram-negative (*S. enterica*), and a spore former (*B. wiedmanii*) previously isolated from milk or the dairy environment to create a mock microbial community that would be inoculated into raw milk. We included an additional Gram-positive bacterium, a member of the *Mycobacteriaceae* family (*M. smegmatis*), because of the potential public health implications and the uniqueness of the cell structure of *Mycobacteria*. Milk samples were specifically inoculated with *Salmonella enterica*, *Listeria monocytogenes*, and *Mycobacterium smegmatis* vegetative cells grown to stationary phase, and *Bacillus wiedmannii* spores. Specific strain information is available in Table 1 and in the Food Microbe Tracker Database (www.foodmicrobetracker.com). Bacterial load of the inoculated milk sample was chosen to represent the largest bacterial concentration allowed for raw milk in the United States, which is 300,000 colony forming units **(CFUs)** per mL, in an attempt to simulate the highest legal bacterial load of incoming milk in a dairy processing plant(33).

**Table 1.** Organisms used in this study.

| Target | Isolate | Source | Accession* |
| --- | --- | --- | --- |
| Total Bacterial DNA/RNA | Pooled Culture |  | NC_000913 |
| *Bacillus wiedmanii* | FSL H7-0344 | Pasteurized 2% milk | NC_004722 |
| *Listeria monocytogenes* | FSL A5-0145 | Raw Milk | NC_003210 |
| *Mycobacterium smegmatis* | FSL X3-0054 | VanDerVen Lab – MC2155 | NC_008596 |
| *Salmonella enterica* | FSL A5-0218 | Raw milk | NC_003197 |
| Bovine DNA | Bovine Blood | Cornell University Dairy Farm | NM_001037471.2 |

\*NCBI accession number of genomes used for primer design; primers for detection of total bacterial DNA target 16S rDNA and were designed based on conserved regions of the 16S gene.

#### Inoculum Preparation

***Bacillus wiedmannii*** spore suspension was prepared according to (34). Briefly, the bacterial isolate was streaked from frozen culture into Brain Heart Infusion (**BHI**) agar (Becton, Dickinson and Co., Sparks, MD) and incubated for 24h at 37°C. Following incubation, a single colony was selected to inoculate a tube containing 5 mL of BHI broth followed by incubation at 37°C for 72h. Next, 100 µL of inoculated BHI was spread plated in duplicate on sporulating media AK Agar #2 (Becton, Dickinson and Co.), which was incubated for 120h at 37°C. Sporulation was confirmed via microscopy with a 7.5% malachite green endospore stain (J.T. Baker, Phillipsburg, NJ) as detailed in (35). Spores were harvested by flooding the agar surface with 10 mL of phosphate-buffered saline (**PBS**) (Weber Scientific, Hamilton, NJ) and scraping the bacterial culture with a cell scraper. Harvested cells were transferred to a sterile centrifuge tube and washed with 10 mL of sterile water three times by centrifugation at 10,500 rpm for 15 minutes and resuspension of the pellet. Following the third wash, 5 mL of sterile water and 5 mL of 100% ethanol (Decon Labs, King of Prussia, PA) were added to the tube and the pellet was resuspended by vortexing. The bacterial pellet resuspended in 50% ethanol was incubated for 12h at 4°C in a rotating platform to eliminate any remaining vegetative cells. After ethanol treatment, the spore suspension was washed another three times, with 10 mL of sterile water as described above. The final spore suspension was kept in sterile water at 4°C until used for spiking experiments.

***Mycobacterium smegmatis*** cells were streaked from frozen stocks into BHI agar, followed by incubation at 37°C for 48h. A single colony was used to inoculate a 5 mL tube containing BHI broth with 1% Tween 80 (Fisher Scientific, Hampton, NH, USA) followed by incubation at 37°C for 48h. The final inoculum was prepared by inoculating 100 µL of liquid culture onto 100 mL of pre-warmed BHI broth with 1% Tween 80, followed by incubation at 37°C for 72h. The resulting stationary phase culture was kept at 4°C until spiking.

***Listeria monocytogenes*** and ***Salmonella enterica*** cells were streaked separately from frozen stocks into BHI agar, followed by incubation at 37°C for 24h. For each strain, a single colony was used to inoculate a 5 mL tube containing BHI broth followed by incubation at 37°C for 24h. The final inoculums were prepared by inoculating 100 µL of each liquid culture onto 100 mL of pre-warmed BHI broth that was subsequently incubated at 37°C for 12h.

At harvesting, bacteria were spiral plated on agar using an Eddy Jet 2W Spiral plater (IUL micro, Barcelona, Spain) at various dilutions to determine bacterial concentrations. Bacterial liquid cultures were kept at 4°C until bacterial enumeration (*B. wiedmanii*, *L. monocytogenes*, and *Salmonella* sp. were kept for 24h and *M. smegmatis* was kept for 48h at 4°C). Inoculum volumes were calculated based on CFUs.

#### Inoculation of Raw Milk

Raw milk was collected from the Cornell University Ruminant Center (CURC – Hartford, NY) bulk tank into sterile 10 oz lock tab containers (Capitol plastics, Amsterdam, NY) and transported on ice to the Milk Quality Improvement Program laboratory in Ithaca, NY. Samples were combined into a single sterile 1000 mL glass bottle and homogenized by inverting the container 50 times. From that bottle, one aliquot (1 mL) was used to make dilutions and determine the initial bacterial count using Standard Plate Count (**SPC**) agar (Milipore Sigma, Burlington, MA), that was incubated for 48h at 32°C. Bacterial enumeration was performed using an automated colony counter (SphereFlash® – Automatic Colony Counter, IUL micro, Barcelona, Spain). A second aliquot (60 mL) was transported on ice to the DairyOne laboratory (DairyOne, Ithaca, NY) for determination of somatic cell counts (**SCC**) and milk contents (fat, protein, lactose, and total solids). The remaining milk was used for inoculation with the strains described above. To allow for interaction between inoculated bacteria and milk components, and to mimic conditions similar to when raw milk is stored in dairy silos, inoculated milk was held at 4°C for 24h prior to use as starting sample for nucleic acid extraction comparisons.

#### Bacterial Enumeration of Inoculated Milk Samples

Total bacterial enumeration was performed through serial dilutions of each milk sample in PBS that was spiral plated in duplicate on standard plate count agar as described above and incubated for 48h at 32°C.

#### Nucleic Acids Extraction

Raw (uninoculated) milk samples were processed in parallel with inoculated milk samples for all kits. Additionally, a no template nucleic acid extraction was carried out as a negative control to assess cross-contamination. As a positive control, a mock bacterial community was created in PBS with the same bacteria described above at the same concentration as the inoculated milk samples; this was included on each extraction plate or run as a control. Inoculated milk samples were extracted in three technical replicates in each of three independent biological replicates, which were performed on separate days with a different raw milk sample used for each biological replicate.

Kits, manufacturers, and protocol details are described in Table 2. For each of the nine protocols evaluated, samples were processed in parallel (uninoculated milk, inoculated milk triplicates, mock bacterial community in PBS, and a negative control without a starting sample “kit buffers only”) according to the manufacturer’s instructions. Exact versions of protocols followed can be found in a GitHub repository as supplementary materials (https://github.com/ErikaGanda/MilkDNA). Extracted nucleic acids were frozen at −80°C until quantification and qPCR assays were performed.

**Table 2.** Nucleic Acid Extraction Kit Characteristics.

| Abbreviations | Kit | Sample Input<br>(μL) | Nucleic Acid Output<br>(μL) | Nucleic Acid | Processing | Capture | Catalog<br>Number | Manufacturer |
| --- | --- | --- | --- | --- | --- | --- | --- | --- |
| PviralDNA<br>PviralRNA | AllPrep PowerViral DNA/RNA kit | 200 | 100 | DNA/RNA | Manual | Column-based | 28000-50 | Qiagen |
| COREDNA<br>CORERNA | MagMAX CORE Nucleic Acid Isolation kit | 200 | 90 | DNA/RNA | Automated | Magnetic-based | A32700 | ThermoFisher |
| ZymoDNA<br>ZymoRNA | ZymoBIOMICs DNA/RNA mini kit | 250 | 100 | DNA/RNA | Manual | Column-based | R2002 | Zymo |
| Mastitis | MagAttract Mastitis | 400 | 100 | DNA | Automated | Magnetic-based | 947757 | Qiagen |
| Pfood | DNeasy PowerFood | 1800 | 100 | DNA | Manual | Column-based | 21000-100 | Qiagen |
| PSoilP | Power Soil Pro | 500 | 100 | DNA | Manual | Column-based | 47014 | Qiagen |
| EZFood | E.Z.N.A. ® Food DNA kit | 500 | 100 | DNA | Manual | Column-based | D4616-00 | Omega |
| RNeasy | RNeasy Protect Bacteria Mini Kit | 1,500 | 50 | RNA | Manual | Column-based | 74524 | Qiagen |
| EZNARNA | E.Z.N.A.® HP Total RNA Kit | 500 | 50 | RNA | Manual | Column-based | R6812-00 | Omega |

#### Nucleic Acids Quantification

Nucleic acid quantification was performed with both a spectrophotometer (Nanodrop 2000, ThermoFisher Scientific) and a fluorescence-based method. Absorbance was measured at 280, 260, and 230 nm for DNA and RNA. Total DNA was also measured with a Quant-iT™ dsDNA High-Sensitivity (HS) Assay Kit (ThermoFisher Scientific); fluorescence measurements were performed using a Synergy H1 plate reader (BioTek Instruments, Winooski, VT, USA) with wavelengths of 490 nm for excitation and 535 nm for emission.

#### qPCR Primer Development and Assay Conditions

For each bacterial strain used to inoculate milk samples, qPCR primers were designed to target the RNA polymerase subunit beta gene (*rpoB*) because it is a single-copy gene and allows for a more accurate comparison between bacterial numbers than 16S rDNA. Primers were also designed to target a conserved region of the 16S gene, and calculated 16S copy numbers were used as a proxy for total bacterial numbers. Primer details are described in Table 3. Reactions were carried out in duplicate using 2 µL of extracted DNA. The final qPCR reaction volumes totaled 25 µL containing 12.5 µL SYBR green master mix (ThermoFisher Scientific), 2 µL extracted DNA, 0.5 µM forward primer, 0.5 µM reverse primer, and 9.5 µL nuclease-free water. Reactions were carried out in a Quantstudio 6 instrument (ThermoFisher Scientific), with the following cycling conditions: 95°C for 10 minutes, followed by 40 cycles of 95°C for 15 seconds and 60°C for 1 minute, and a melting curve: 95°C for 15 seconds, 60°C for 1 minute, and 95°C for 15 seconds.

**Table 3.**
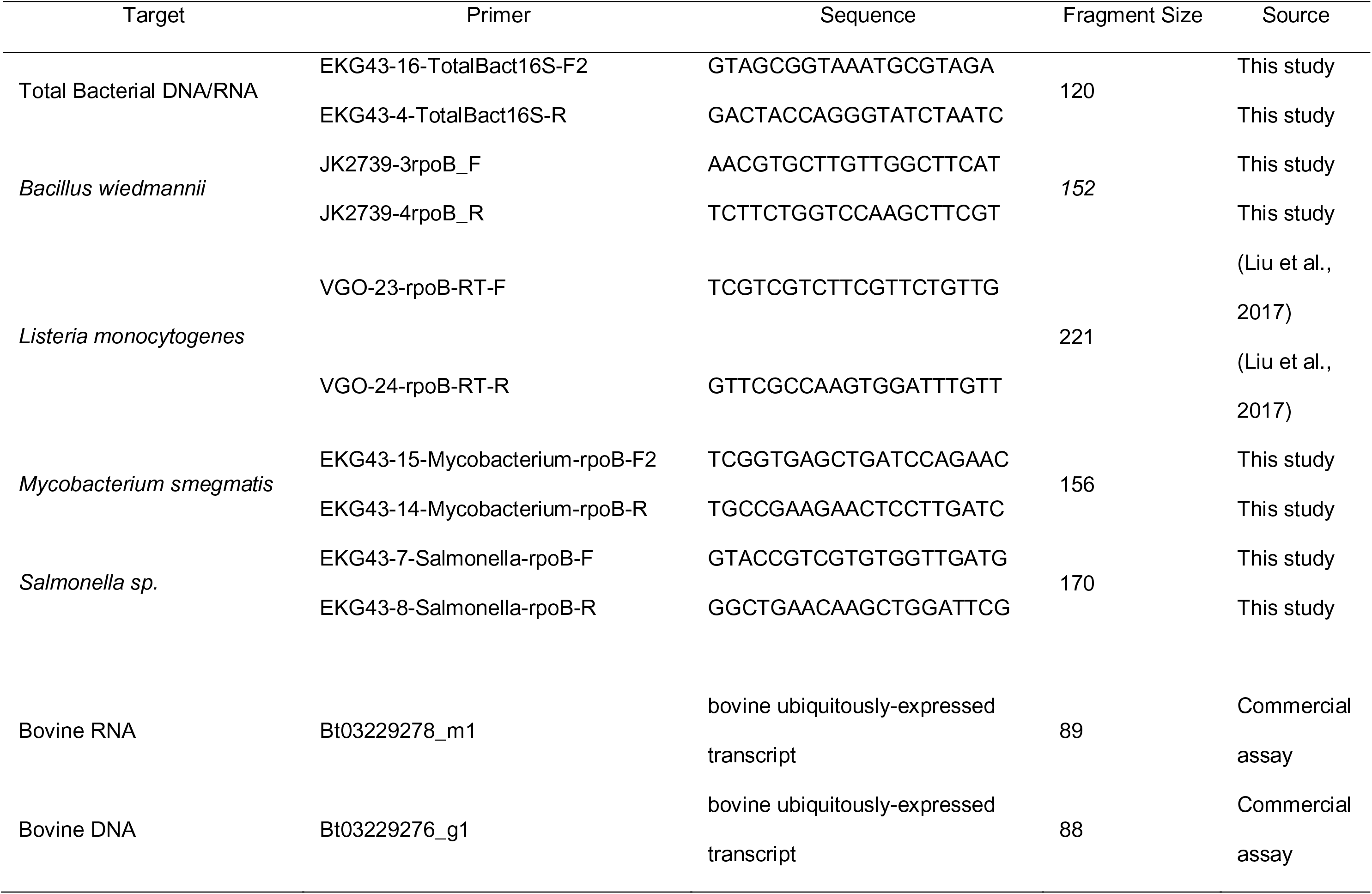
Primers Used in this Study.

Bovine genome copy numbers were calculated using a commercial TaqMan assay targeting the UXT gene (ThermoFisher Scientific). The final qPCR reaction totaled 20 µL, and included 1 µL of gene expression assay, 10 µL TaqMan Fast Advanced Master Mix (ThermoFisher Scientific), 7 µL of nuclease-free water, and 2 µL of template. Reactions were carried out in a Quantstudio 6 instrument, with the following cycling conditions: 95°C for 20 seconds, followed by 40 cycles of 95°C for 1 second and 60°C for 20 seconds. For each of the three biological replicates, samples extracted with all kits were amplified in a single PCR plate and compared using the same standard curve.

#### qPCR Data Analysis

Amplification data were exported from QuantStudio Real-Time PCR Software (ThermoFisher Scientific) into Excel (version 16.0.11325.20156, Microsoft Corp., Redmond, WA). Standard curves were built with serial dilutions of purified bacterial DNA that was quantified using a Nanodrop spectrophotometer (ThermoFisher) and genome equivalents in each reaction were calculated as described in (36). Standard curves with an R2 < 0.9 and efficiency < 70% were discarded and reactions were repeated. For 2 out of 24 reaction plates, one outlier was removed based on visual inspection of deviating standard curve data points.

Copies per µL of DNA input were calculated for each reaction. Melt curves were visually inspected and cycle threshold (**CT**) values of samples in which melt curves did not match standard samples’ peaks were not used for final copy number calculations. Because different kits required varying amounts of sample input and nucleic acid output (Table 2), data was normalized to allow for comparison between kit protocols. We chose to simulate copy numbers in 1 mL of milk input and 100 µL DNA output for each kit, using the following equations:

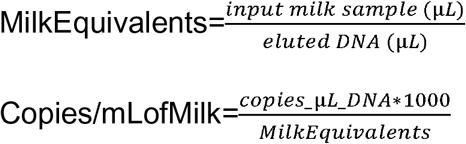

Normalized copy number data were log-transformed prior to statistical analysis. The final database was cross-referenced and checked with the open-source software openrefine (https://github.com/OpenRefine). All statistical analyses with inoculated milk PCR data were performed in R (version 3.4.3; R project, Vienna, Austria).

#### Statistical Model Selection

Linear models were fit using the “lm” function in R to compare log copy numbers between kits. For each assay, three linear models were compared; including (i) experiment number (SpikeSet) only, (ii) including qPCR efficiency only, and (iii) both (i) and (ii) as covariates. Given that only three experiments were performed, experiment number was treated as a fixed effect rather than a random effect. To account for different variances observed across kits, models not assuming homoscedasticity were also fit using generalized least-squares using the nlme (37) package in R. Models were inspected by Q-Q plots and residual vs. predicted plots. The model with best fit as determined by smallest AIC was used as the final model. Estimated marginal means of each model were calculated using the emmeans (38) package and are depicted in graphs overlaid to scatterplots of original datapoints. Raw data and code are available on a GitHub repository: https://github.com/ErikaGanda/MilkDNA.

### Host Depletion Protocols

#### Selective Osmotic Lysis of Host Cells and Host DNA Depletion through Propidium Monoazide (PMA) Treatment of Uninoculated Raw Milk

An osmotic lysis-based host DNA depletion protocol was adapted from (27). Briefly, 500 µL of uninoculated milk were centrifuged at 10,000g for 8 minutes, whey was discarded while fat and pellet were kept in the microcentrifuge tube. Five hundred microliters of sterile double distilled water (**ddH_2_O**) were added to the pellet. The tube was vortexed until pellet dissolution followed by incubation at room temperature for 5 minutes to allow for osmotic lysis of mammalian cells. Four different concentrations of PMA (Biotium, Hayward, CA, catalog # 40019) were compared: 10, 20, 40, and 50µM. Untreated milk (milk that was not centrifuged or exposed to osmotic lysis) and milk exposed to osmotic lysis but not exposed to PMA were also included in comparisons. To account for biological variation and assess repeatability, experiments were performed in three biological replicates performed on three different days, using a different raw milk sample for each biological replicate.

After incubation at room temperature, the appropriate volume of PMA was added to each tube to achieve desired concentrations, followed by briefly mixing, and incubation in the dark (in an aluminum-foil wrapped box) for 5 minutes on a rotating platform.

Following PMA incubation in the dark, PMA was inactivated by light exposure. Samples were placed horizontally on ice, < 20 cm from a 500W halogen light source (Woods Halogen Work Light, Southwire Company LLC, Carrollton, GS, USA). Samples were exposed to light on a rotating platform for 5 minutes, and frozen at −80°C until DNA extraction with a magnetic-based method (MagMAX CORE Nucleic Acid Purification Kit – ThermoFisher). Quantitative PCR reactions and data analysis were performed as described above, with the exception that only bovine DNA and total bacterial copy numbers were quantified, as no bacteria was inoculated in the milk samples used. Linear models were fitted to assess the effect of selective lysis in non-PMA treated samples, (NTC versus 0µM) and the effect of PMA concentration on copy numbers.

#### Selective Enzymatic Lysis of Host Cells and Host DNA depletion through PMA Treatment in Inoculated Raw Milk

Because milk has more fat, protein, and minerals then saliva we hypothesized that the 1:1 osmotic lysis included in the protocol adapted from (27) was not optimal for lysing bovine cells in milk compared to human cells in saliva.

We thus also evaluated a combination of a mild lysis solution followed by incubation with two concentrations of subtilisin, a protease from *Bacillus licheniformis* (Krackeler Scientific, cat# 45-P5380-25MG). The lysis solution contained 7.6 g/L sodium carbonate, 8.8 g/L sodium bicarbonate, 2.43 g/L disodium EDTA, and 2.71 g/L tetrasodium EDTA. Reagents were solubilized in 1L of sterile double distilled water using a stir plate with a magnetic rotating bar for 6 hours, prior to pH measurement and titration to 9.5, and filter-sterilization with a 0.22 µm filter (Corning Disposable Vacuum Filter – Fisher Scientific, Waltham, MA).

Experiments were performed in two separate days with a newly collected milk sample in each day. To access potential lysis biases, we spiked milk with a Gram-positive and a Gram-negative bacterium as described above (*L. monocytogenes* and *Salmonella* sp.). Spiked milk samples were treated prior to DNA extraction as follows: Four hundred microliters of inoculated milk were added to 1.6 mL of lysis solution and incubated at room temperature for 5 minutes prior to the addition of subtilisin. Two concentrations of subtilisin were tested: 10 mg/mL (10x) and 1 mg/mL (1x) – to achieve final concentrations of 0.73 and 0.073 units per reaction, respectively. After adding the enzyme, samples were incubated at 50°C for 5 minutes and placed on ice after incubation. Samples were then centrifuged at 10,000g for 8 minutes. The supernatant containing lysis solution and enzyme was discarded, and the pellet was resuspended with 400µL of PBS. Treated samples were exposed to PMA at 20 µM as described above, photo-activated using two light sources (Halogen lamp described above or BLU-V System – Qiagen) and stored at −80°C until DNA extraction with a magnetic-based method (MagMAX CORE extraction kit – ThermoFisher).

Quantitative PCR reactions and data analysis were performed as described above, with the exception that Bovine DNA, Total bacterial copy numbers, *L. monocytogenes*, and *Salmonella* were quantified, as we hypothesized that the lysis solution could affect Gram-negative bacteria differently than Gram-positive bacteria.

#### Comparison of Host DNA Depletion Methods in Uninoculated Raw Milk

Based on initial experiments with various PMA concentrations we decided to use 20µM to include in a comparison with three commercial host DNA depletion kits. Kits, manufacturers, and protocol details are described in Table 4. Samples were processed according to the manufacturer’s instructions and frozen at −80°C.

**Table 4.**
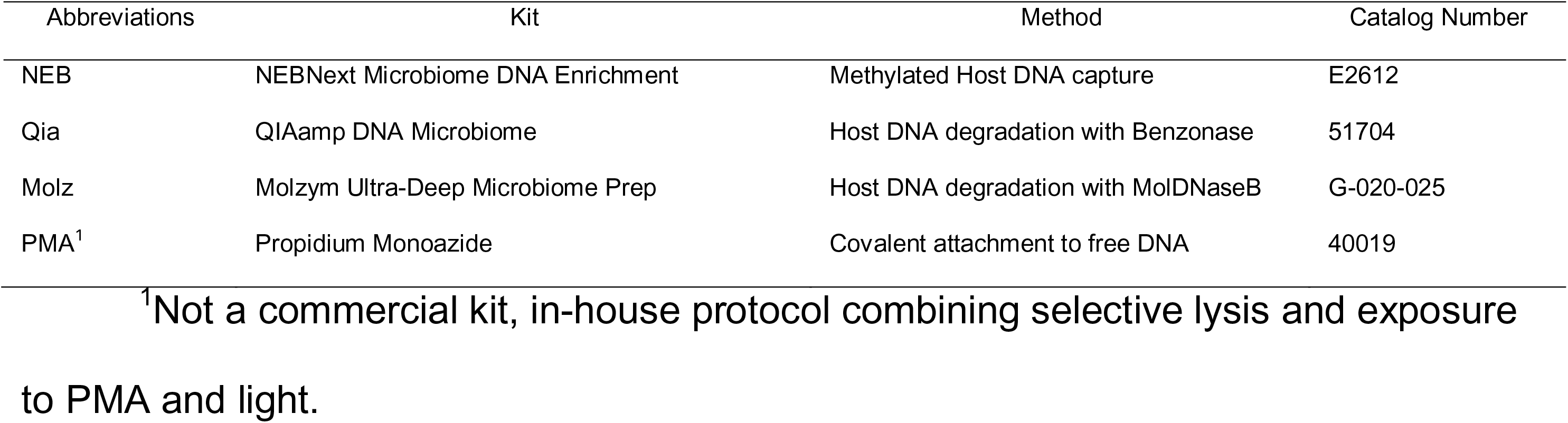
Host DNA depletion Kit Characteristics.

For the NEB host depletion method, DNA was extracted with a magnetic-based extraction and quantified using a qubit fluorometer (ThermoFisher) prior to methylated host DNA capture reactions. Final microbial enriched DNA cleanup was performed using AMPure magnetic beads (Beckman Coulter). Exact versions of protocols followed can be found in our GitHub repository https://github.com/ErikaGanda/MilkDNA. Three biological replicates (each with two technical replicates) were prepared using a new sample of raw milk for each biological replicate (with the exception of MolZym kit, which was only performed with duplicate milk samples in one day due to limitation in available reagent amounts).

Quantitative PCR reactions and data analysis were performed as described above, with the exception that only bovine DNA and total bacterial copy numbers were quantified, as no bacteria was inoculated in the milk samples used.

#### Sequencing

In addition to qPCR, we performed deep untargeted sequencing of a subset of nine samples at the Cornell Biotechnology Resource Center. Briefly, DNA quality control was performed in a fragment analyzer and sequencing libraries were constructed using a Nextera DNA Flex kit (now renamed to Illumina DNA Prep, Illumina, San Diego, CA) and 2×150 BP paired-end sequencing was performed in an Illumina NextSeq500. Sequencing data quality control and in silico host signal removal was performed as previously described (3, 4), and microbial data was processed as described in Beck et al. 2020 (4). Briefly, adapter removal and quality trimming was performed with TrimGalore (39) and trimmed reads of at least 50bp in length were classified using Kraken v0.3 (40) against a multi-eukaryotic database of 31 common food ingredients and contaminants including the bovine reference genome (GCF_000003205.7, Btau_5.0.1) for bioinformatic removal of the host signal. Then taxonomic profiling of samples was completed using Kraken v0.3 against microbial RefSeq genomes as described in Beck et. al., 2020 (4). Principal Component Analysis (PCA) and Robust PCA (41) were performed in pseudocounted Isometric Log-Ratio transformed microbial reads (4).

#### Data availability

Raw data and code are available on our GitHub repository: https://github.com/ErikaGanda/MilkDNA. Raw sequencing reads have been deposited into SRA under the submission number SUB7938697.

## RESULTS

### Extraction Method Significantly Impacts Bacterial Quantification through qPCR

We assessed seven commercially available DNA extraction methods on their ability to isolate DNA from milk samples inoculated with a mock bacterial community (for milk sample characteristics see Supplementary Table S1). Nucleic acid quantification and quality measurements (Supplementary Table S2) need to be interpreted with caution due to the low overall DNA yield (<10 ng/µL for all samples), which placed readings below the linear range for most samples. However, all kits yielded sufficient DNA to allow for qPCR-based quantification of the members of the bacterial mock community (i.e., *Bacillus wiedmanii*, *Listeria monocytogenes*, *Mycobacterium smegmatis*, and *Salmonella* sp.) as well as total bacterial 16S rDNA copies and bovine DNA. Quantitative PCR data (Figure 1) demonstrated that the two magnetic-based DNA extraction methods (COREDNA and Mastitis, the two leftmost methods in Fig. 1) always yielded numerically higher copy numbers (for all members of the mock community and for total bacterial 16S rDNA) than the other five kits; these values were generally significantly higher (except for *L. monocytogenes*, where values for one non-magnetic kit did not differ significantly from the two magnetic kits). Bacterial copy numbers for the two magnetic-based kits also had lower variability between technical replicates than column-based kits, with E.Z.N.A. Food DNA kit showing particularly large variability between technical replicates; DNA yield for this kit were also so low that no qPCR amplification was observed in the *L. monocytogenes* qPCR targeting *rpoB*.

**Figure 1.**
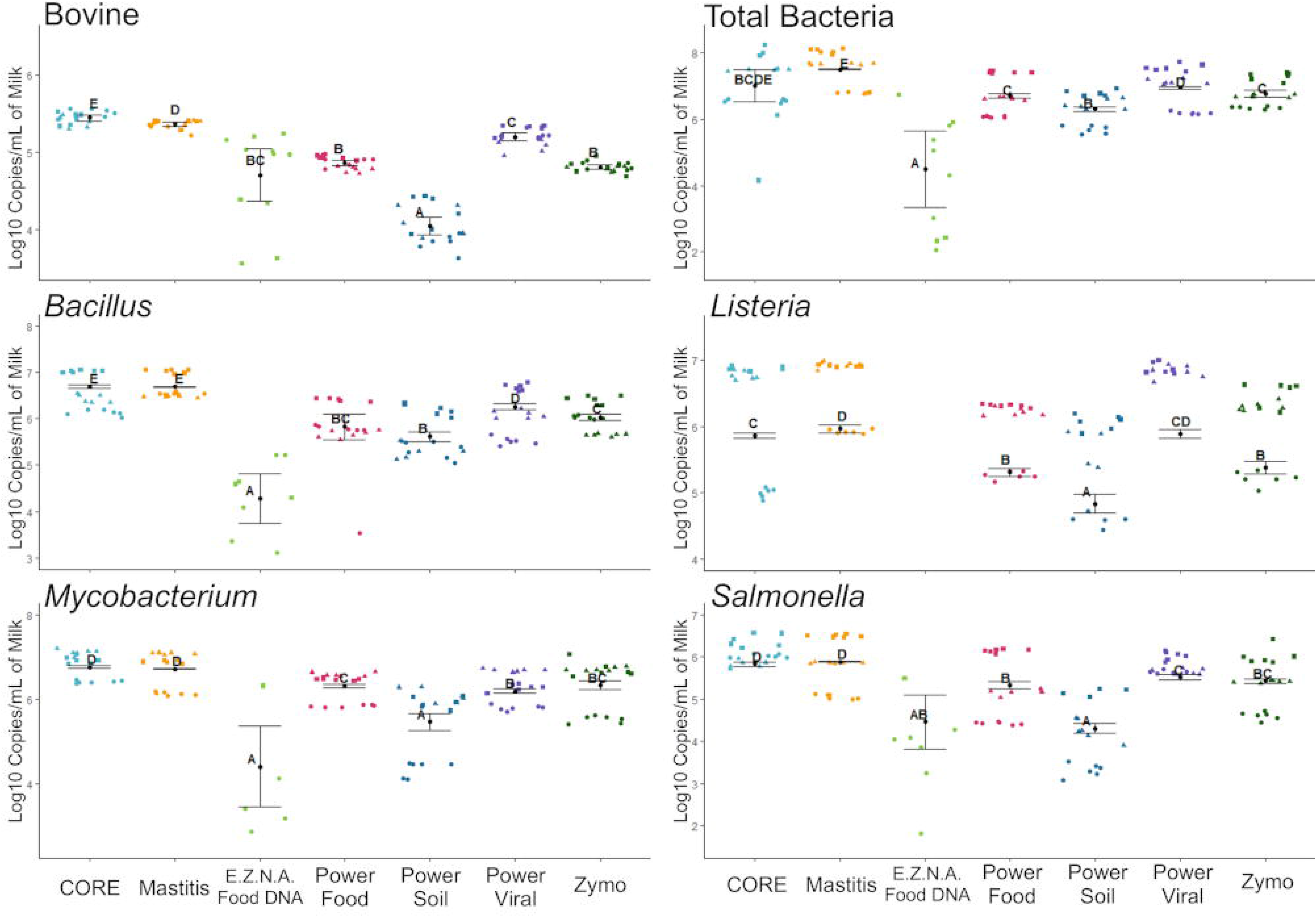
Scatterplots of raw data of normalized log copy numbers per mL of milk obtained with different DNA extraction kits. Bars overlaid represent model estimated marginal means and standard deviations. Different letters represent significantly different means at the 0.05 level after Tukey multiple comparison adjustment. Scatterplot shapes represent each of three independent biological replicates (circles - first; triangles - second; and squares - third., whereas colors represent kits evaluated. Each panel depicts results of a single gene target evaluated.

Among the column-based extraction methods, Power Food, Power Viral, and Zymo were comparable with regard to (i) their ability to recover DNA of the members of the Mock community and (ii) variability within technical replicates (see Fig. 1; Table 5 provides detailed descriptive statistics). The Power Soil kit (another column-based kit), generally provided lower bacterial DNA yields for each member of the mock community than these three kits, even though the differences in log10 copies per ml of milk were only significantly for some organisms (e.g., *Mycobacterium*).

**Table 5.** Descriptive Statistics of Log Copies per mL of Milk – DNA extraction.

| Organism | Mean log10 copies per mL of milk as detected via qPCR (StdDev), N |  |  |  |  |  |  |
| --- | --- | --- | --- | --- | --- | --- | --- |
|  | COREDNA | Mastitis | EZFood | PFood | PSoilP | PviralDNA | ZymoDNA |
| Total Bacterial DNA | 6.99 (0.96),18 | 7.51 (0.54),18 | 4.30 (1.72),12 | 6.71 (0.57),18 | 6.31 (0.48),18 | 6.96 (0.59),18 | 6.78 (0.40),18 |
| *Bacillus wiedmanii* | 6.53 (0.39),18 | 6.67 (0.25),18 | 4.31 (0.77),12 | 5.79 (0.67),18 | 5.61 (0.45),18 | 6.09 (0.49),18 | 6.02 (0.32),18 |
| *Listeria monocytogenes* | 6.21 (0.89),18 | 6.58 (0.48),18 | NA (NA)<sup>a</sup>,12 | 5.93 (0.49),18 | 5.46 (0.68),18 | 6.84 (0.09),18 | 6.00 (0.59),18 |
| *Mycobacterium smegmatis* | 6.81 (0.31),18 | 6.70 (0.42),18 | 3.97 (1.39),12 | 6.29 (0.34),18 | 5.44 (0.82),18 | 6.25 (0.38),18 | 6.30 (0.58),18 |
| *Salmonella sp.* | 6.07 (0.26),18 | 5.80 (0.61),18 | 3.83 (1.12),12 | 5.24 (0.72),18 | 4.13 (0.77),18 | 5.77 (0.19),18 | 5.34 (0.61),18 |
| Bovine DNA | 5.45 (0.08),18 | 5.37 (0.05),18 | 4.71 (0.59),12 | 4.86 (0.08),18 | 4.05 (0.24),18 | 5.20 (0.11),18 | 4.81 (0.06),18 |

<sup>a</sup> Not applicable, this method did not yield detectable signal in qPCR for *Listeria monocytogenes*.

We also evaluated the ability to isolate bacterial RNA for (i) two kits designed for isolation of RNA only and for (ii) two kits designed for isolation of both RNA and DNA (see Table 2). Similar to the DNA detection data detailed above, all kits yielded very low total nucleic acid concentrations based on spectrophotometry. Reverse transcriptase qPCR amplification of gene targets for *Salmonella* sp. and *L. monocytogenes* as well as bovine RNA did not yield amplification on samples treated with DNase (these initial tests were performed on two technical replicates for each of the four kits). Control amplifications on nucleic acids before DNAse treatment however, yielded amplification with CT values that did not differ between the RT-qPCR and the control qPCR targeting DNA, suggesting the presence of residual DNA in the extracted nucleic acids.

### Osmotic Lysis followed by PMA Treatment Decreases Host DNA but also Impacts Bacterial DNA

We compared protocols that combined osmotic lysis and treatment with multiple concentrations of PMA for their ability to reduce host DNA, compared to bacterial DNA copy numbers. To assess if osmotic lysis alone impacted copy-numbers, we compared non-treated control (NTC) samples to samples that underwent centrifugation and osmotic lysis with ddH_2_O, but no PMA addition (0µM PMA); this comparison revealed significantly lower bovine and bacterial DNA copy numbers in 0µM PMA samples, suggesting significant effects of sample processing (including lysis) independent of PMA addition (Figure 2, NTC vs. 0µM comparisons). We also observed that PMA concentration significantly impacted bovine and bacterial DNA copy numbers in a dose-dependent manner: as PMA concentration increased, a sharp decrease was observed in bovine copy-numbers, whereas a less pronounced, but still noticeable decrease was observed in bacterial copy-numbers (Figure 2). This trend was confirmed in the model estimates, which showed a significant and negative effect of PMA in both linear models, with a greater estimate for bovine copy numbers (−0.022) versus bacterial copy numbers (−0.012). Treatment with 20 µM PMA was the concentration deemed optimal as it yielded the greatest decrease in host DNA without critically compromising bacterial DNA recovery, and thus was chosen for subsequent comparisons with commercially available host DNA depletion protocols.

**Figure 2.**
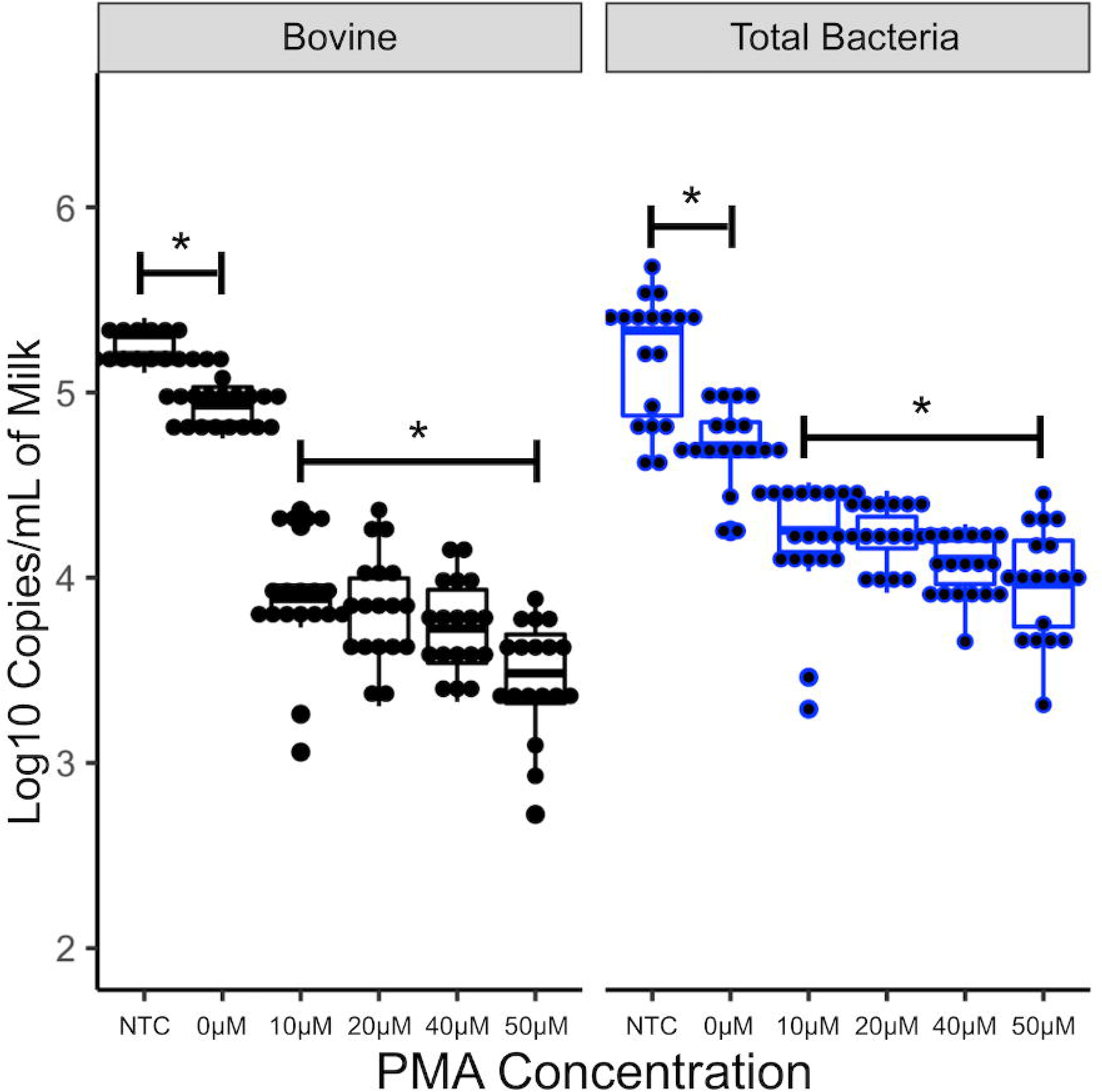
Effect of osmotic lysis of raw milk followed by treatment with various PMA concentrations on host and bacterial DNA counts as determined by qPCR. Boxplots represent normalized log copy numbers per mL of milk. NTC: untreated sample, 0µM: milk that underwent osmotic lysis but was not treated with PMA. Asterisks represent significant differences at p < 0.05 in linear model comparisons. Osmotic lysis significantly decreases Log10 copies in non-PMA treated samples (NTC vs 0µM) and increasing PMA concentrations significantly decreases Log10 Copies in a dose-dependent manner.

We also selected a subset of milk samples from each of the three biological replicates to undergo deep untargeted sequencing, including one sample each of the three biological replicates representing (i) untreated samples, (ii) samples treated with 20 µM PMA and (iii) samples that underwent osmotic lysis but were not treated with PMA (underwent centrifugation, selective lysis with ddH_2_O but were not treated with PMA); (iii) was included to allow us to determine if differences observed would be due to PMA treatment or sample processing prior to PMA addition.

Sequencing had an average read depth of 51,563,707 reads per sample (range 25,488,728 to 127,203,413). Overall less than 1% of all reads that passed quality control were determined to be of microbial origin regardless of PMA treatment (Table 6, Figure 3), and around 3% remained unclassified.

**Table 6.**
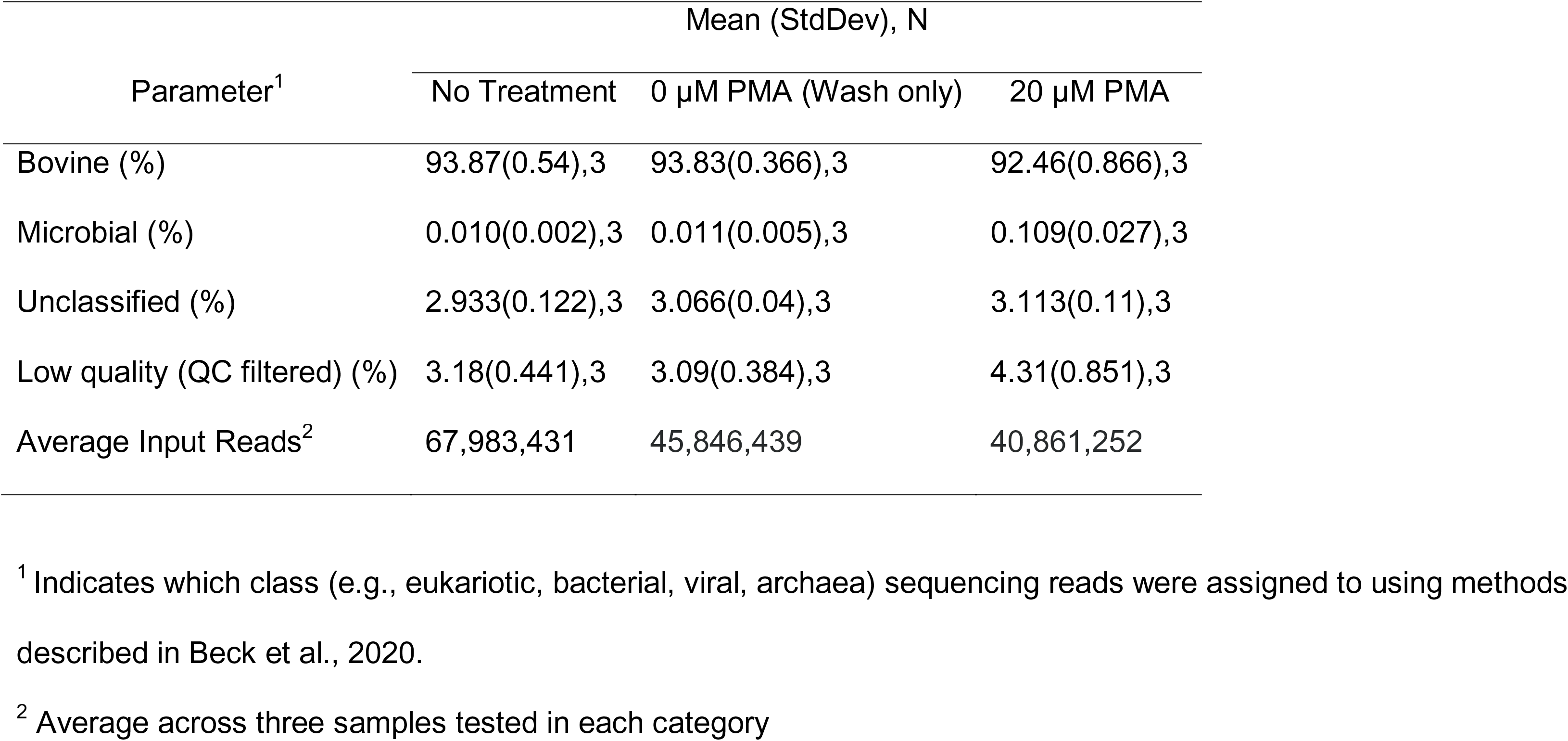
Descriptive statistics of shotgun metagenomics sequencing results assessing the effect of PMA treatment.

| Parameter<sup>1</sup> | Mean (StdDev), N |  |  |
| --- | --- | --- | --- |
|  | No Treatment | 0 µM PMA (Wash only) | 20 µM PMA |
| Bovine (%) | 93.87(0.54),3 | 93.83(0.366),3 | 92.46(0.866),3 |
| Microbial (%) | 0.010(0.002),3 | 0.011(0.005),3 | 0.109(0.027),3 |
| Unclassified (%) | 2.933(0.122),3 | 3.066(0.04),3 | 3.113(0.11),3 |
| Low quality (QC filtered) (%) | 3.18(0.441),3 | 3.09(0.384),3 | 4.31(0.851),3 |
| Average Input Reads<sup>2</sup> | 67,983,431 | 45,846,439 | 40,861,252 |

<sup>1</sup> Indicates which class (e.g., eukariotic, bacterial, viral, archaea) sequencing reads were assigned to using methods described in Beck et al., 2020. <sup>2</sup> Average across three samples tested in each category

**Figure 3.**
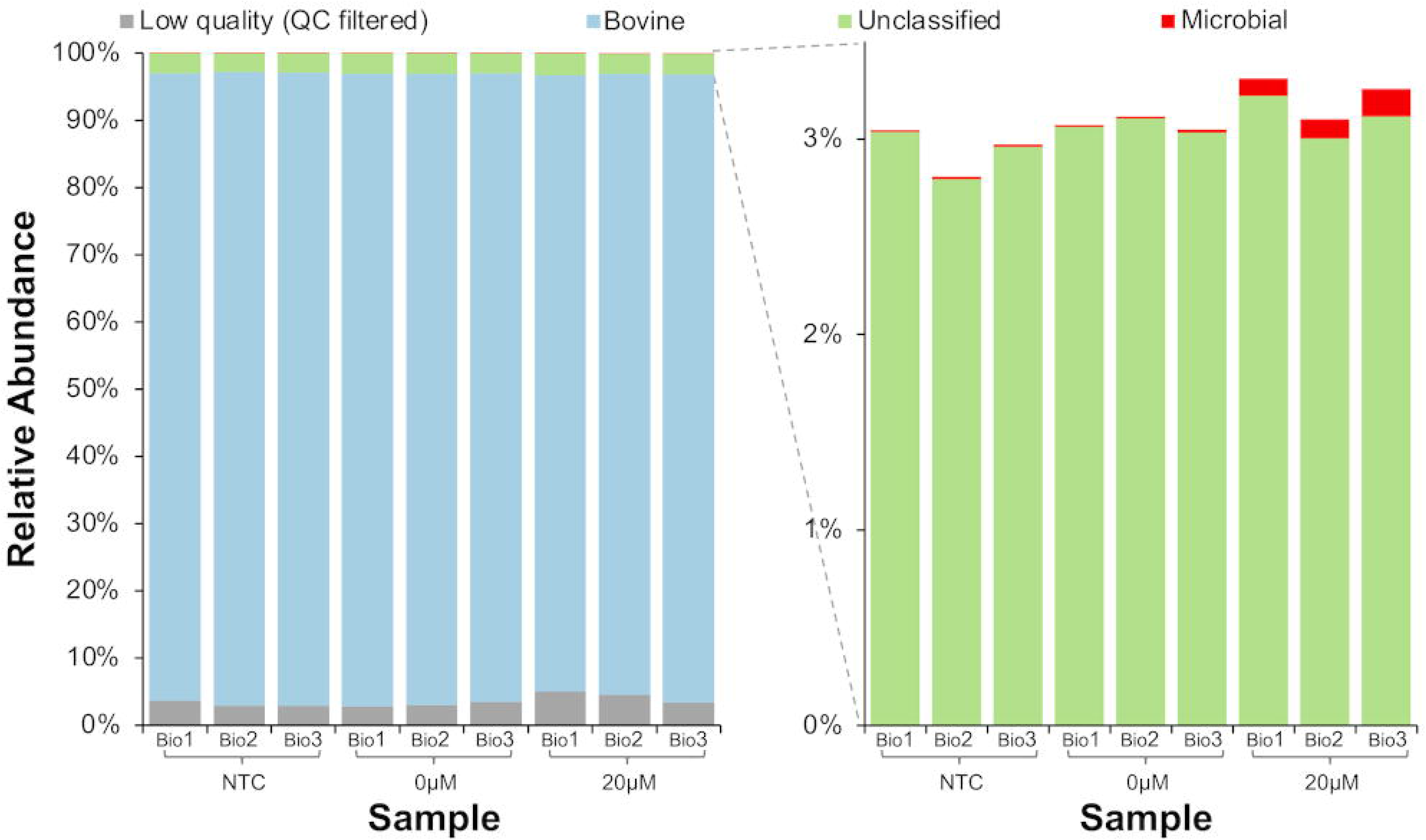
Effect of host DNA depletion with PMA on percentage of sequencing reads assigned to the bovine genome or microbial genomes. Raw milk was collected in three separate days and divided into aliquots that underwent each of the processing methods, including (i) NTC: untreated sample, (ii) 0µM: milk that underwent osmotic lysis but was not treated with PMA, and (iii) 20µM: milk that underwent osmotic lysis and PMA treatment at 20 µM.

While we observed a pronounced increase in the number of bacterial reads per one million sequenced reads with the PMA treatment (Figure 4A), attempting to deplete host DNA with PMA also appeared to influence the resulting microbial profiles of samples, leading to visible differences between replicates of the same biological samples (Figure 4B). For example, PMA treatment of Bio1 sample resulted in greater relative abundances of *Cutibacterium* when compared to aliquots of the same sample (Bio1) that were not treated (NTC) or underwent selective lysis with ddH_2_O but no PMA treatment (0µM). Similar differences between NTC, 0µM, and 20µM were observed for the other two samples sequenced (Bio2 and Bio3, Figure 4B).

**Figure 4.**
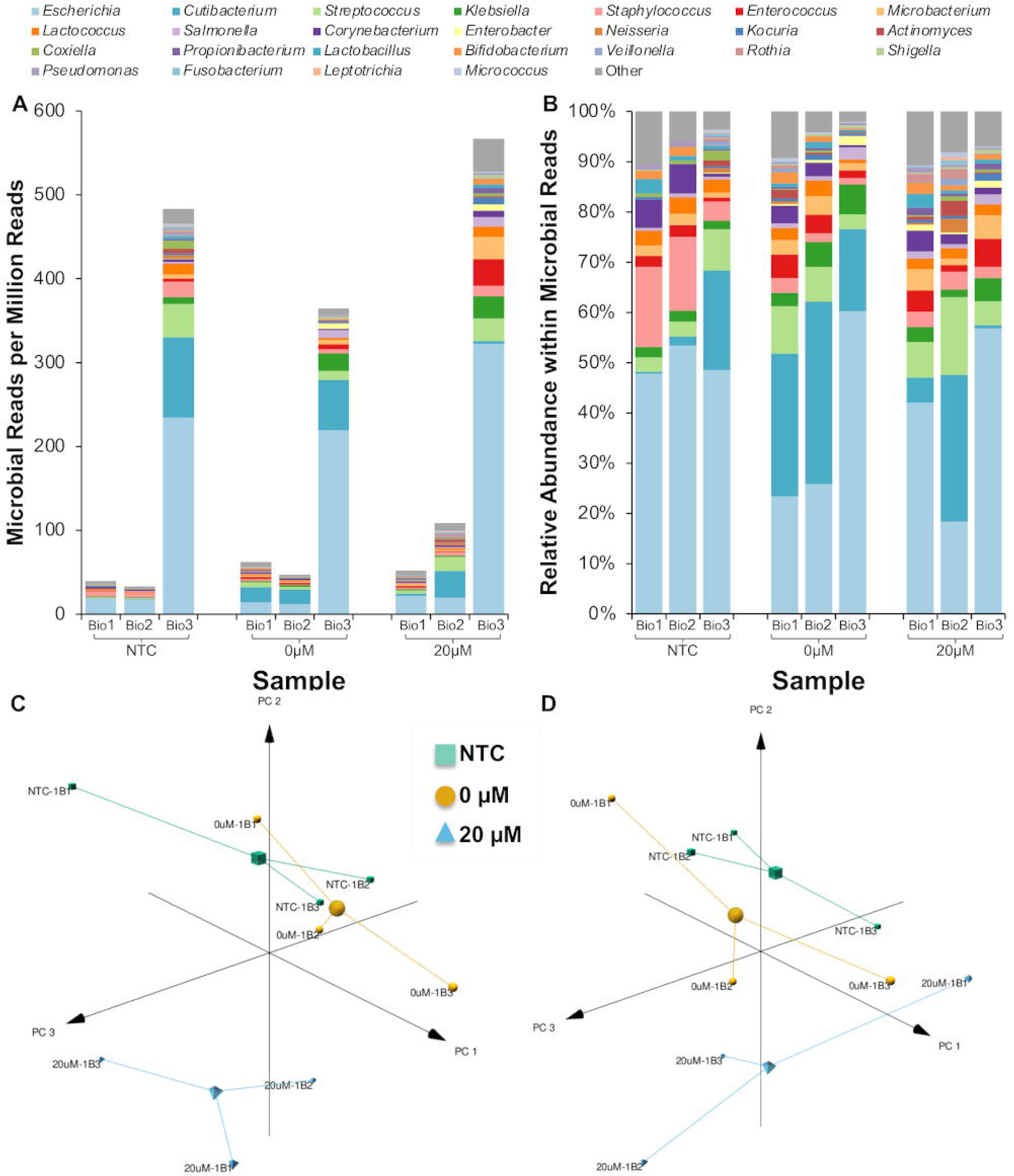
Microbial profile of samples exposed to osmotic lysis and PMA treatment. (A) Microbial reads per million reads by biological sample and treatment. (B) Relative abundance within microbial reads by biological sample and treatment. (C) Principal Component Analysis (PCA) and (D) Robust PCA performed in Isometric Log-Ratio transformed microbial reads. NTC: untreated sample, 0µM: milk that underwent osmotic lysis but was not treated with PMA, 20µM: milk that underwent osmotic lysis and PMA treatment at 20 µM.

The effect of PMA treatment was also seen when the first 3 principal components were plotted in PCA (Figure 4C) or robust PCA (Figure 4D). We observed that PMA treated samples separated from other samples (all have negative PC_2) instead of samples clustering by biological sample origin regardless of PMA treatment. Robust PCA(42) was used in an effort to alleviate the effects of PMA treatment; however, it was not able to effectively improve the signal-to-noise ratio introduced by PMA treatment.

### Host Depletion Methods Successfully Decrease Host DNA, But Not to the Extent Needed to Make a Significant Impact for Use in Milk Sequencing Studies

We compared host DNA depletion methods and osmotic lysis followed by PMA treatment in three independent biological replicates. In each biological replicate, a freshly collected milk sample was homogenized and divided into five separate aliquots that were processed in parallel for each of the five methods (e.g., PMA method, Molzym kit, NEB kit, Qiagen kit, and negative controls [NTC]). Means by which host DNA depletion was accomplished varied between methods: some kits depended on host DNA degradation followed by treatment with different enzymes (e.g., Qiagen and Molzym kits), while the NEB kit was based on the capture of methylated host DNA (for details on kits and methods, see Table 4, for descriptive statistics, see Table 7). Overall, host DNA removal treatments significantly decreased bovine copy numbers when compared to a negative control (Figure 5A). However, we also observed a significant loss of bacterial copy numbers across methods (Figure 5B). Notably, the Molzym kit caused the largest reduction of both bovine and bacterial DNA copy numbers to levels that likely would be insufficient as inputs for shotgun metagenomics sequencing. Treatment of raw milk with Molzym reagents specifically resulted in a pellet that was challenging to bring back to solution, which likely led to considerable nucleic acid losses and renders this method unreliable for use with milk.

**Table 7.**
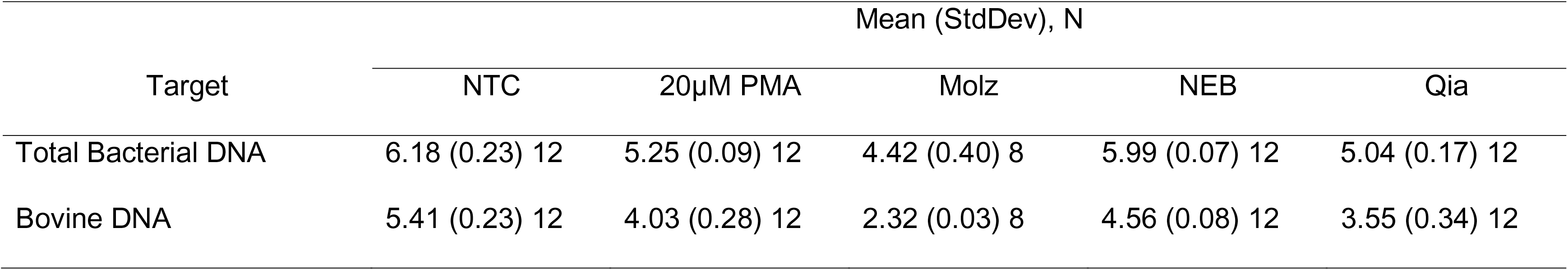
Descriptive Statistics of Log Copies per mL of Milk – Host Depletion.

| Target | Mean (StdDev), N |  |  |  |  |
| --- | --- | --- | --- | --- | --- |
|  | NTC | 20µM PMA | Molz | NEB | Qia |
| Total Bacterial DNA | 6.18 (0.23) 12 | 5.25 (0.09) 12 | 4.42 (0.40) 8 | 5.99 (0.07) 12 | 5.04 (0.17) 12 |
| Bovine DNA | 5.41 (0.23) 12 | 4.03 (0.28) 12 | 2.32 (0.03) 8 | 4.56 (0.08) 12 | 3.55 (0.34) 12 |

**Figure 5.**
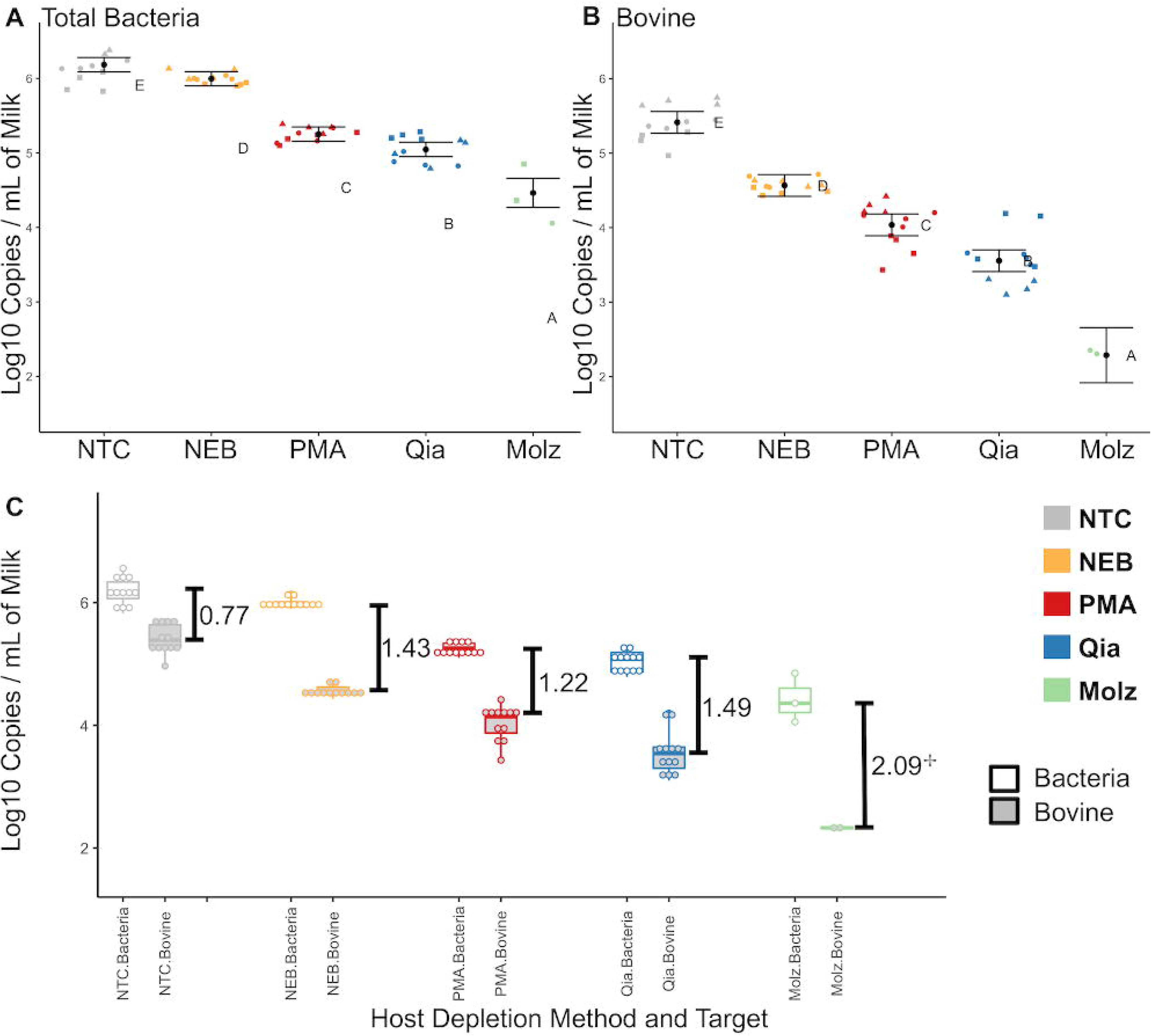
Comparisons of host depletion methods in uninoculated milk. Scatterplots of raw data of normalized log copy numbers per mL of milk. Bars overlaid represent model estimated marginal means and standard deviations for comparisons performed on Total Bacterial copy numbers (A) and Bovine copy numbers (B) and. Different letters represent significant Tukey-adjusted differences at the 0.05 level. Scatterplot shapes represent each independent experiment, whereas colors represent kits evaluated. Paired boxplots (C) represent normalized log copy numbers per mL of milk of two independent experiments with two technical replicates each, with exception of Molzym, that only had technical replicates performed in one day. Numbers represent the log difference between mean log Bacterial copy numbers and mean log Bovine copy numbers. ^[]^Denotes that technical replicates were only performed in one day due to sample becoming an insoluble pellet during processing.

As all host DNA depletion methods also led to the depletion of bacterial DNA, we choose to use the calculated log difference between bovine and bacterial DNA copy numbers as key metrics for evaluating the different host DNA depletion methods. Untreated (NTC) samples showed an average of 0.77 log higher bacterial DNA copy number as compared to bovine DNA copy numbers (Figure 5C). By comparison, the log difference after different host depletion methods ranged from 1.22 log (PMA method) to 2.09 log (Molzym kit). If treated samples underwent untargeted sequencing, this decrease in bovine copies would translate into negligible differences in terms of relative abundances of bacterial reads versus bovine reads given the approximately thousand-fold difference between the bovine genome size and average bacterial genome sizes.

### Enzymatic Selective Lysis of Inoculated Milk followed by PMA Treatment Differentially Affects Detection of Gram-positive and Gram-negative Bacteria through qPCR

In a final attempt to optimize a host depletion protocol that would be applicable to raw milk, we tested a lysis protocol that included a mild protease, with the objective of more efficiently permeabilizing mammalian cells while keeping bacterial cells intact. Because we were aware of potential biases that could be introduced by adding an enzymatic lysis step prior to PMA treatment, we decided to inoculate the samples tested with Gram-positive and Gram-negative bacteria to address potential differential bacterial permeabilization by subtilisin that would result in DNA inactivation by PMA binding. We also investigated if the light source would have an effect on the efficiency of PMA binding by processing samples in parallel and exposing PMA-treated sample duplicates to either a halogen-light source or a commercial apparatus designed for use with PMA treated samples (BLU-V, Qiagen). We prepared two biological replicates with two technical replicates for each comparison, from which duplicate qPCR reactions were done.

Treatment with subtilisin decreased the number of culturable bacteria as assessed through CFU plate counts of milk that had been treated with lysis solution prior to PMA exposure (Figure 6A). However, we did not observe a difference in CFU in milk samples treated with different enzyme concentrations. The light source did not affect copy numbers (*P=0.74*, data not shown); therefore qPCR comparisons were performed in combined data from both halogen light source and BLU-V apparatus for Figure 6B. As expected, we observed a decrease in bovine copy numbers in subtilisin-PMA treated samples compared to negative control to a much greater extent than what we observed with osmotic lysis (Figure 6B, first panel). However, we also observed a dose-dependent decrease in Gram-negative copy numbers in treated samples (Figure 6B, third panel), while no differences were observed in Gram-positive copy numbers (Figure 6B, second panel). This trend was confirmed by a less steep but still noticeable decrease in total bacterial copy numbers when compared to Gram-negative copy numbers (Figure 6B, fourth panel). These data led us to conclude that enzymatic treatment followed by PMA inactivation differentially affects Gram-negative bacteria copy numbers, leading to biased results if used to prepare DNA for sequencing purposes.

**Figure 6.**
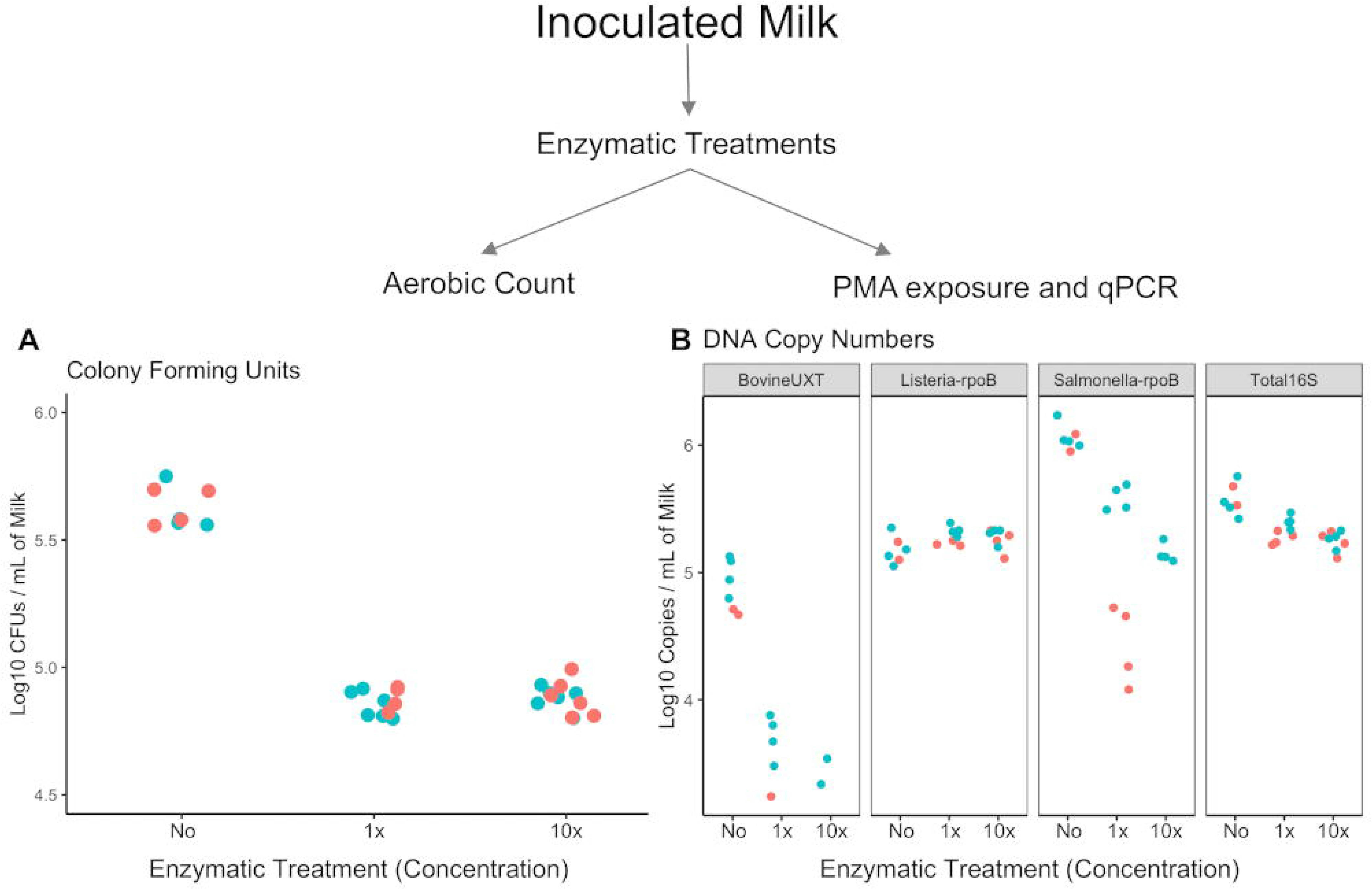
Comparison of enzymatic lysis prior to host DNA depletion with 20 µM of PMA inoculated milk. Aerobic standard plate count results (A) and qPCR results (B). Different colors correspond to two independent experiments with two technical replicates each, from with duplicate qPCR reactions were done. CT values of samples in which melt curves did not match standard samples’ peaks were not used.

## DISCUSSION

In this study we comprehensively evaluated both nucleic acid extraction and host depletion protocols in bovine raw milk. The rationale of using raw milk as a model included the recent publication of several studies highlighting the variability of raw milk microbiota (43–45); the potential influence of raw milk microbial quality on processed dairy products (46–48); and the fact that patterns of the milk microbiome represent potential “biomarkers” that could be tracked (19, 20). All of these studies highlight milk as a potential candidate for using HTS as part of quality assurance and risk assessment in the food industry in the future. Our results indicate that (i) magnetic-based DNA extraction methods are superior for extracting DNA from milk, (ii) although host DNA depletion methods decrease the amount of bovine DNA in a given sample, the reduction would not be sufficient for effectively depleting bovine DNA in HTS studies of raw milk, and (iii) we observed biases introduced by certain protocols; both on the overall microbial profile as well as selective bias against Gram-negative bacteria.

### Magnetic-based DNA extraction is superior to column-based methods for raw milk samples

We compared several nucleic acid extraction protocols on bovine raw milk. While all protocols evaluated for extraction of DNA were able to successfully extract (albeit low amounts of) total DNA, we observed higher variability in replicates in some protocols when compared to others. The low DNA concentration from milk extracts is in agreement with previous reports (24, 25, 49). One particular aspect of this study is that we inoculated different types of bacteria that would be of interest in a dairy processing environment into raw milk and performed targeted qPCR to assess differential extraction of Gram-positive and Gram-negative bacteria. While most protocols were able to successfully extract DNA from the bacteria inoculated, magnetic-based DNA extraction approaches had the best recovery and the lowest inter-replicate variability as evaluated by qPCR. It must be noted that the differences observed in the recovery of bacterial DNA are a characteristic of processing steps rather than a flaw or inefficiency of a given protocol (i.e. while some extraction methods carry the entire sample lysate from one step to the following, some protocols require fixed or maximum volumes to be transferred to the next step due to volume limitations in reaction tubes used). It is important to highlight that different DNA extraction methods have advantages and drawbacks, which can also vary according to the material they are applied to. As no general gold-standard method exists to date for this application, the selection of an extraction method should be based on the objectives of each particular study (50). Nevertheless, a growing consensus exists on the importance of mechanical disruption such as bead-beating for microbiome applications (51), and magnetic-based extractions have been demonstrated to be particularly effective in the diagnosis of tuberculosis in serum and plasma of patients (52) and the extraction of algae DNA for NGS studies (53), highlighting magnetic methods as a common theme around diverse nucleic-acid based applications.

### Extracting bacterial RNA from high-quality raw milk is challenging

The methods used here co-purify both DNA and RNA, or each nucleic acid separately, and based on inoculation of bacterial loads in some cases exceeding regulatory standards for raw milk; we expected that bacterial RNA should be detectable. We observed very low concentrations of extracted DNA and were unable to successfully detect bacterial RNA from our inoculated milk samples. This was a surprising finding, as literature on the milk metatranscriptome is available (18). However, the protocols previously used to successfully isolate RNA from milk involved extensive centrifugation of large sample volumes followed by several wash steps (54), as well as samples with much greater bacterial loads, including dairy products such as cheese, which contain high levels of organisms responsible for fermentation of the products (18, 20).

### Current methods for host DNA depletion are not suitable for application in untargeted HTS studies of milk

Because the bovine cells are present in milk along with the microbes of interest, and due to the striking difference between bovine and bacterial genome sizes, developing efficient yet unbiased host-DNA depletion methods is critical for the adoption of HTS technologies in food safety. While we observed a decrease in the number of bovine copies when compared to bacterial copies as assessed through qPCR in all protocols evaluated, this decrease would not be sufficient to effectively change the relative abundance of reads being assigned to the bovine genome in HTS studies. This was confirmed when we performed deep untargeted sequencing in a subset of samples that had demonstrated promising results in qPCR but still detected over 99% of reads mapping to the bovine genome independent of host DNA depletion.

Among the other host DNA depletion protocols evaluated were immunoprecipitation of methylated eukaryotic DNA and selective lysis of mammalian cells followed by DNAse treatment. One reason for the lack of efficiency observed in the treatment of milk with either of these methods could be the challenges associated with extracting DNA from milk in the first place. Such protocols require large amounts of good quality DNA, which is practically impossible to obtain from milk samples without extensive centrifugation and pellet washes, potentially proving impractical in industry settings and representing added opportunities for sample contamination or unintentional bias of the microbiota for lipid- or protein-bound microorganisms. The characteristics of milk in itself can also pose a challenge (presence of fats, proteins, and ions) (8, 9). Challenges specific to nucleic acid extraction from milk have been discussed in the literature to some extent, specifically Metzger et al. reported challenges in amplifying bacterial DNA from DNA extracted from milk (55).

To the best of our knowledge, this was the first study to evaluate host DNA depletion methods in a food matrix. Limited literature is available trying to address host contamination in clinical sequencing applications for pathogen detection, with reported improvements in the detection of malaria (56) and pathogens in infected tissue samples (57) and sputum (31). Reports are also available describing attempts of using host DNA depletion in cerebrospinal (29), and arthroplasty (28) fluids, as well as human saliva (27). The common theme around these studies was the varying efficiencies across methods and sample types, highlighting the need for individual assessment of host depletion methods using the desired sample type.

### Enzymatic-based selective lysis followed by PMA treatment differentially affects Gram-negative bacteria in inoculated raw milk

Propidium monoazide has been extensively used to differentiate between live and dead bacterial cells due to its inability to penetrate intact cell membranes and has recently been indicated to be an effective method for host DNA depletion for use in HTS studies (27, 58, 59). We thus developed an enzymatic-based method for host DNA depletion (which utilized a mild protease) combined with subsequent PMA treatment in an effort to better lyse mammalian cells while retaining intact bacterial cells.

While we detected an effect of the enzymatic treatment on total bacterial numbers as measured through plate counts that appeared to be independent of enzymatic concentration, we did not observe a decrease in Gram-positive bacterial numbers as measured through qPCR after PMA exposure. Nevertheless, we observed an enzyme dose-dependent decrease of Gram-negative copy numbers in samples that underwent selective lysis followed by PMA exposure. These data led us to conclude that enzymatic treatment might have differentially permeabilized the membrane of Gram-negative cells, allowing PMA to bind to DNA without observable differences in overall bacterial survival. Taken together, these observations suggest that potential biases could occur against Gram-negative bacteria detection through qPCR in this method.

### It is imperative to optimize protocols for samples with different characteristics

The generalizability of this study lies on the importance of standardization and validation of methods for each specific food matrix. It is also important to highlight the challenges associated with low bacterial biomass samples and the fact that various foods have a wide range of biomass, from very low in raw milk, to relatively high microbial content in some cultured dairy products. Efforts to standardize and recommend best practices in HTS studies have recently begun, particularly pertaining to low biomass samples (60–62) and must be continued.

## CONCLUSION

Our results suggest that magnetic-based extraction methods are superior for bacterial nucleic acid isolation from bovine milk. Host DNA remains a challenge for untargeted sequencing of milk, highlighting that the food matrix characteristics should always be considered whenever planning HTS studies. Enzymatic-lysis based PMA host depletion introduced dose-dependent biases against Gram-negative bacteria, suggesting that selective lysis permeabilized Gram-negative to PMA, which subsequently hindered our ability to detect Gram-negative bacteria through qPCR without affecting live bacterial counts. While it is not possible to test all methods available at any given time, we focused on kits and protocols that have been most widely used for the extraction of nucleic acids from milk. As procedures are improved, or new methods are developed, reevaluation of protocols available would prove useful, as the development of HTS-based tools to aid and improve quality assurance and food safety programs continue to hold great promise.

## DECLARATIONS

### Ethics Approval

All experimental procedures were carried out at Cornell University, because no animal sampling was necessary for this study, an approved research protocol by the Cornell University Institutional Animal Care and Use Committee was not necessary.

### Consent for publication

Not applicable.

### Availability of data and material

The datasets and R code used and/or analyzed during the current study are available on https://github.com/ErikaGanda/MilkDNA. Raw sequencing reads have been deposited into SRA under the submission number SUB7938697.

### Authors’ contributions

Conceived and designed the experiments: **EKG, MW** Performed the experiments: **EKG** Performed automated DNA extraction: **BC, RRA** Analyzed the data: **EKG, KLB, NH, BK** Wrote the paper: **EKG** Revised the manuscript: **EKG, KLB, NH, BK, LG, MW**. All authors reviewed the manuscript.

## Acknowledgements

We thank members of CURC for allowing us to sample, Dr. VanDerVen Lab for providing us *Mycobacterium smegmatis* MC2-155 strain, and Dr. Mann Lab for providing bovine RNA used as a standard.

---

**Supplementary Table S1.**
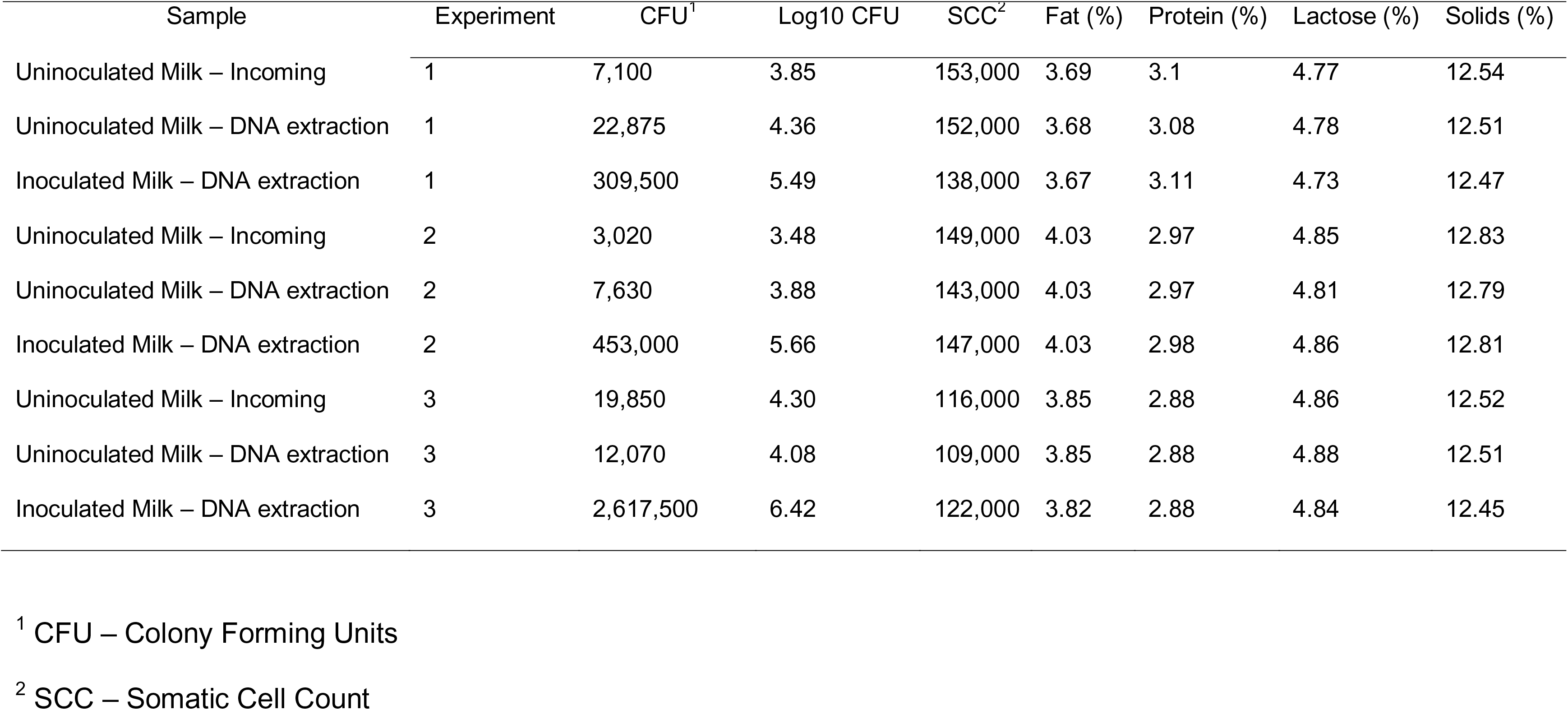
Milk Sample Characteristics.

**Supplementary Table S2.**
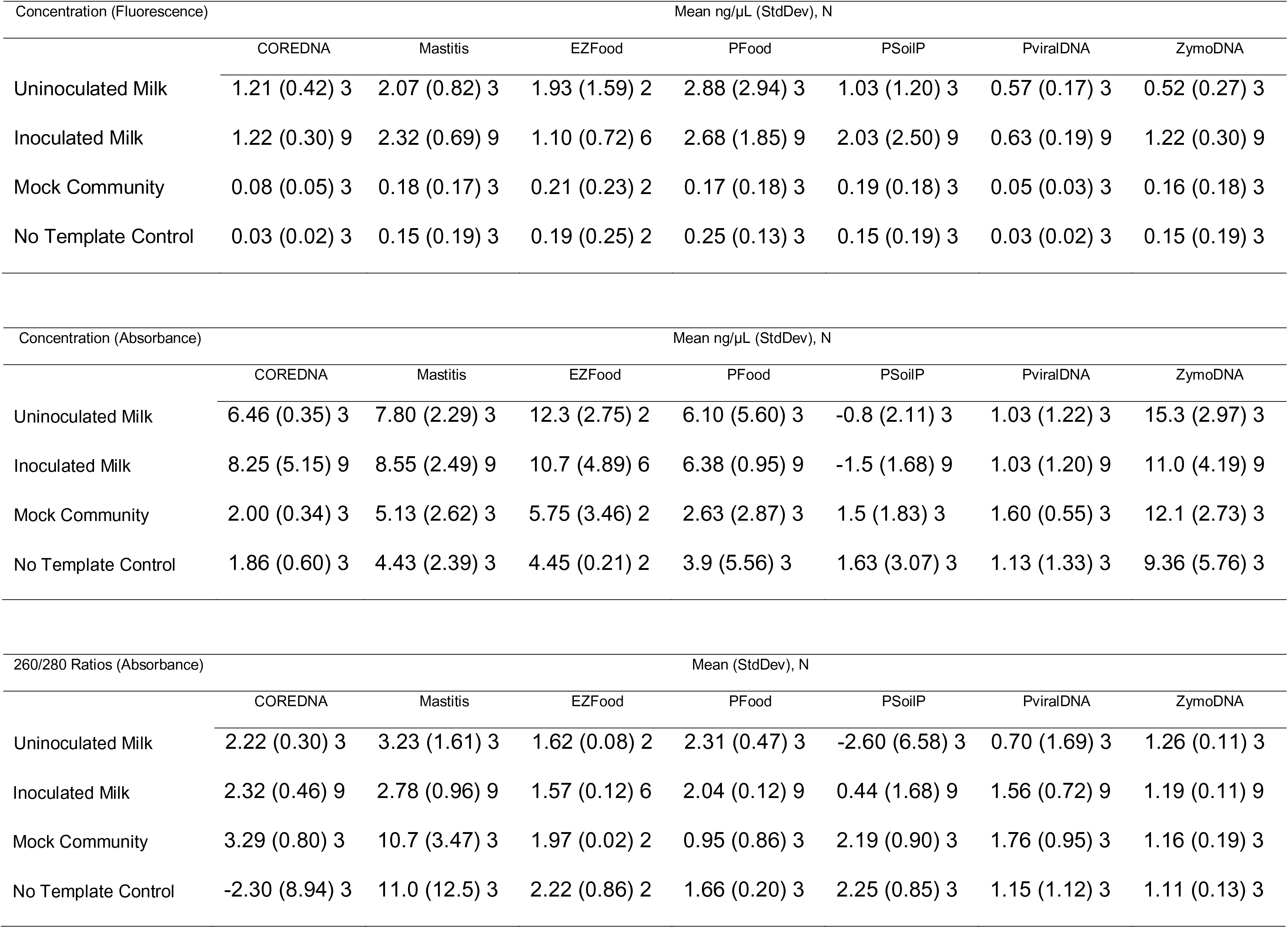
Descriptive Statistics of Nucleic Acid Quantification and Quality Measurements.

