## Supplemental Table S1 for "DNA extraction and host depletion methods significantly impact and potentially bias bacterial detection in a biological fluid"

**Supplementary Table S1 – Milk Sample Characteristics**

| Sample | Experiment | CFU^1^ | Log10 CFU | SCC^2^ | Fat (%) | Protein (%) | Lactose (%) | Solids (%) |
| --- | --- | --- | --- | --- | --- | --- | --- | --- |
| Uninoculated Milk – Incoming | 1 | 7,100 | 3.85 | 153,000 | 3.69 | 3.1 | 4.77 | 12.54 |
| Uninoculated Milk – DNA extraction | 1 | 22,875 | 4.36 | 152,000 | 3.68 | 3.08 | 4.78 | 12.51 |
| Inoculated Milk – DNA extraction | 1 | 309,500 | 5.49 | 138,000 | 3.67 | 3.11 | 4.73 | 12.47 |
| Uninoculated Milk – Incoming | 2 | 3,020 | 3.48 | 149,000 | 4.03 | 2.97 | 4.85 | 12.83 |
| Uninoculated Milk – DNA extraction | 2 | 7,630 | 3.88 | 143,000 | 4.03 | 2.97 | 4.81 | 12.79 |
| Inoculated Milk – DNA extraction | 2 | 453,000 | 5.66 | 147,000 | 4.03 | 2.98 | 4.86 | 12.81 |
| Uninoculated Milk – Incoming | 3 | 19,850 | 4.30 | 116,000 | 3.85 | 2.88 | 4.86 | 12.52 |
| Uninoculated Milk – DNA extraction | 3 | 12,070 | 4.08 | 109,000 | 3.85 | 2.88 | 4.88 | 12.51 |
| Inoculated Milk – DNA extraction | 3 | 2,617,500 | 6.42 | 122,000 | 3.82 | 2.88 | 4.84 | 12.45 |

^1^ CFU – Colony Forming Units

^2^ SCC – Somatic Cell Count
