## Supplemental Table S2 for "DNA extraction and host depletion methods significantly impact and potentially bias bacterial detection in a biological fluid"

**Supplementary Table S2 – Descriptive Statistics of Nucleic Acid Quantification and Quality Measurements**

| Concentration (Fluorescence) | Mean ng/µL (StdDev), N | | | | | | |
| --- | --- | --- | --- | --- | --- | --- | --- |
|  | COREDNA | Mastitis | EZFood | PFood | PSoilP | PviralDNA | ZymoDNA |
| Uninoculated Milk | 1.21 (0.42) 3 | 2.07 (0.82) 3 | 1.93 (1.59) 2 | 2.88 (2.94) 3 | 1.03 (1.20) 3 | 0.57 (0.17) 3 | 0.52 (0.27) 3 |
| Inoculated Milk | 1.22 (0.30) 9 | 2.32 (0.69) 9 | 1.10 (0.72) 6 | 2.68 (1.85) 9 | 2.03 (2.50) 9 | 0.63 (0.19) 9 | 1.22 (0.30) 9 |
| Mock Community | 0.08 (0.05) 3 | 0.18 (0.17) 3 | 0.21 (0.23) 2 | 0.17 (0.18) 3 | 0.19 (0.18) 3 | 0.05 (0.03) 3 | 0.16 (0.18) 3 |
| No Template Control | 0.03 (0.02) 3 | 0.15 (0.19) 3 | 0.19 (0.25) 2 | 0.25 (0.13) 3 | 0.15 (0.19) 3 | 0.03 (0.02) 3 | 0.15 (0.19) 3 |

| Concentration (Absorbance) | Mean ng/µL (StdDev), N | | | | | | |
| --- | --- | --- | --- | --- | --- | --- | --- |
|  | COREDNA | Mastitis | EZFood | PFood | PSoilP | PviralDNA | ZymoDNA |
| Uninoculated Milk | 6.46 (0.35) 3 | 7.80 (2.29) 3 | 12.3 (2.75) 2 | 6.10 (5.60) 3 | -0.8 (2.11) 3 | 1.03 (1.22) 3 | 15.3 (2.97) 3 |
| Inoculated Milk | 8.25 (5.15) 9 | 8.55 (2.49) 9 | 10.7 (4.89) 6 | 6.38 (0.95) 9 | -1.5 (1.68) 9 | 1.03 (1.20) 9 | 11.0 (4.19) 9 |
| Mock Community | 2.00 (0.34) 3 | 5.13 (2.62) 3 | 5.75 (3.46) 2 | 2.63 (2.87) 3 | 1.5 (1.83) 3 | 1.60 (0.55) 3 | 12.1 (2.73) 3 |
| No Template Control | 1.86 (0.60) 3 | 4.43 (2.39) 3 | 4.45 (0.21) 2 | 3.9 (5.56) 3 | 1.63 (3.07) 3 | 1.13 (1.33) 3 | 9.36 (5.76) 3 |

| 260/280 Ratios (Absorbance) | Mean (StdDev), N | | | | | | |
| --- | --- | --- | --- | --- | --- | --- | --- |
|  | COREDNA | Mastitis | EZFood | PFood | PSoilP | PviralDNA | ZymoDNA |
| Uninoculated Milk | 2.22 (0.30) 3 | 3.23 (1.61) 3 | 1.62 (0.08) 2 | 2.31 (0.47) 3 | -2.60 (6.58) 3 | 0.70 (1.69) 3 | 1.26 (0.11) 3 |
| Inoculated Milk | 2.32 (0.46) 9 | 2.78 (0.96) 9 | 1.57 (0.12) 6 | 2.04 (0.12) 9 | 0.44 (1.68) 9 | 1.56 (0.72) 9 | 1.19 (0.11) 9 |
| Mock Community | 3.29 (0.80) 3 | 10.7 (3.47) 3 | 1.97 (0.02) 2 | 0.95 (0.86) 3 | 2.19 (0.90) 3 | 1.76 (0.95) 3 | 1.16 (0.19) 3 |
| No Template Control | -2.30 (8.94) 3 | 11.0 (12.5) 3 | 2.22 (0.86) 2 | 1.66 (0.20) 3 | 2.25 (0.85) 3 | 1.15 (1.12) 3 | 1.11 (0.13) 3 |
