## Supplemental Table S3 for "DNA extraction and host depletion methods significantly impact and potentially bias bacterial detection in a biological fluid"

**Supplementary Table S4 – Detailed description of qPCR results for Bovine, Total Bacteria, and Bacillus.**

|  |  | **Bovine DNA** | | | | **Total Bacterial DNA** | | | | ***Bacillus*** | | | |
| --- | --- | --- | --- | --- | --- | --- | --- | --- | --- | --- | --- | --- | --- |
| **Replicate** | **Extraction Kit** | **Mean** | **StDev** | **Failures** | **Total** | **Mean** | **StDev** | **Failures** | **Total** | **Mean** | **StDev** | **Failures** | **Total** |
| First | COREDNA | 5.51 | 0.05 | 4 | 12 | 6.11 | 0.63 | 2 | 12 | 6.13 | 0.06 | 6 | 12 |
| First | EZFood | 5.10 | 0.15 | 4 | 12 | 4.36 | 1.26 | 2 | 12 | 4.16 | 0.82 | 5 | 12 |
| First | Mastitis | 5.30 | 0.09 | 4 | 12 | 5.84 | 1.36 | 0 | 12 | 5.93 | 1.13 | 4 | 12 |
| First | Pfood | 4.80 | 0.15 | 4 | 12 | 5.30 | 1.37 | 0 | 12 | 5.27 | 0.82 | 4 | 12 |
| First | PSoilP | 3.83 | 0.10 | 4 | 12 | 5.14 | 1.00 | 0 | 12 | 5.16 | 0.32 | 4 | 12 |
| First | PviralDNA | 5.24 | 0.05 | 4 | 12 | 5.63 | 0.86 | 2 | 12 | 5.52 | 0.09 | 6 | 12 |
| First | ZymoDNA | 4.85 | 0.06 | 4 | 12 | 5.88 | 0.78 | 0 | 12 | 5.70 | 0.62 | 4 | 12 |
| Second | COREDNA | 5.39 | 0.07 | 4 | 12 | 6.33 | 2.02 | 0 | 12 | 5.46 | 1.92 | 2 | 12 |
| Second | Mastitis | 5.40 | 0.02 | 4 | 12 | 6.18 | 2.18 | 0 | 12 | 6.01 | 0.86 | 4 | 12 |
| Second | Pfood | 4.81 | 0.08 | 4 | 12 | 5.70 | 1.50 | 0 | 12 | 4.43 | 1.97 | 2 | 12 |
| Second | PSoilP | 4.06 | 0.17 | 4 | 12 | 5.24 | 2.00 | 0 | 12 | 5.29 | 0.28 | 4 | 12 |
| Second | PviralDNA | 5.10 | 0.08 | 4 | 12 | 6.14 | 1.68 | 1 | 12 | 5.91 | 0.46 | 4 | 12 |
| Second | ZymoDNA | 4.83 | 0.09 | 4 | 12 | 5.73 | 1.47 | 0 | 12 | 5.45 | 0.36 | 4 | 12 |
| Third | COREDNA | 5.47 | 0.10 | 4 | 12 | 6.27 | 1.86 | 4 | 12 | 6.83 | 0.35 | 4 | 12 |
| Third | EZFood | 4.56 | 0.69 | 4 | 12 | 4.37 | 2.08 | 6 | 12 | 4.50 | 0.19 | 9 | 12 |
| Third | Mastitis | 5.38 | 0.06 | 4 | 12 | 7.35 | 0.94 | 2 | 12 | 6.67 | 0.64 | 4 | 12 |
| Third | Pfood | 4.91 | 0.06 | 4 | 12 | 6.37 | 1.76 | 1 | 12 | 6.30 | 0.17 | 5 | 12 |
| Third | PSoilP | 4.26 | 0.19 | 4 | 12 | 5.98 | 1.36 | 1 | 12 | 6.08 | 0.23 | 4 | 12 |
| Third | PviralDNA | 5.26 | 0.08 | 4 | 12 | 6.65 | 1.47 | 1 | 12 | 6.42 | 0.48 | 4 | 12 |
| Third | ZymoDNA | 4.84 | 0.09 | 4 | 12 | 6.18 | 1.59 | 0 | 12 | 5.97 | 0.79 | 4 | 12 |

^*^ Ezfood was not included in comparisons with the second biological replicate because it was backordered at the time.
